# Simple and Multiplexed Enrichment of Rare DNA Variants via Sequence-Selective and Temperature-Robust Amplification

**DOI:** 10.1101/136465

**Authors:** Lucia R. Wu, Sherry X. Chen, Yalei Wu, Abhijit A. Patel, David Yu Zhang

## Abstract

Rare DNA sequence variants hold important clinical and biological information, but are challenging for existing methods (e.g. PCR, NGS) to profile in an inexpensive, multiplexed, simple-to-implement, and sequence-general way. Here, we present Blocker Displacement Amplification (BDA), a temperature-robust PCR method that selectively amplifies all sequence variants within a roughly 20 nt window by 1000-fold over wildtype sequences, allowing easy detection and quantitation of hundreds of potentials variants originally at ≤0.1% allele frequency. BDA employs a rationally designed competitive hybridization reaction to achieve similar enrichment performance across anneal temperatures ranging from 56°C to 64°C. This temperature robustness facilitates multiplexed enrichment of many different variants across the genome, and furthermore enables the use of inexpensive and portable thermocycling instruments for rare DNA variant detection. To show the sequence generality of BDA, we demonstrated enrichment on 156 single-nucleotide variants (SNVs). BDA has been validated on multiple different PCR platforms, DNA polymerases, and sample types including clinical cell-free DNA samples collected from the blood plasma of lung cancer patients. BDA quantitation of mutation allele fraction is generally consistent with deep sequencing results.

---

The economical and high-throughput detection and quantitation of nucleic acid sequence variants is a key goal in the road to widespread adoption of precision medicine, wherein optimal individualized treatment is provided to each patient based on his or her unique genetic and disease profile. Profiling rare nucleic acid variants with low allele frequencies, such as cancer mutations in cell-free DNA [1–3] or drug resistance in pathogen sub-populations [4–6], presents a challenge for current molecular diagnostic technologies [7]. Allele-specific PCR methods [8–12] are difficult to scale to allow highly multiplexed rare variant detection, and deep sequencing approaches [13–15] are not economical because they waste a large majority of their read capacity on sequencing wildtype templates and amplicons. Sample enrichment to elevate the allele fractions of rare variants can allow economical sequencing-based rare variant profiling, but has been difficult to realize in highly multiplexed settings.

Past demonstrations of DNA variant enrichment employ either selective depletion of wildtype sequences via hybridization [16–19] or selective PCR amplification of variants [20– 23]. It has been challenging to scale these approach to multiplexed enrichment of many different variants across different loci, because existing methods require that the operational reaction temperature sits in the “Goldilocks” zone of every single blocker or probe. To elaborate, a wildtype sequence at a particular locus may bind to its probe or blocker to form a duplex with melting temperature *T*_*M*_, and a variant at that locus would form a duplex with melting temperature (*T*_*M*_ − Δ*T*_*M*_); only if the reaction temperature is in the range between these two values is there significant enrichment. Despite more than 40 years of biophysical studies, the melting temperature of a duplex sequence can only be predicted with a standard error of roughly 1.4*°*C [25, 26], corresponding to a 95% confidence interval that spans 5.6*°*C. Empirical optimization is impractical in highly multiplexed systems, due to the combinatorially many interactions possible among DNA species that each could influence *T*_*M*_.

Here, we present a simple method to enrich hundreds of potential DNA sequence variants from multiple genomic loci simultaneously. Simple here denotes both that little to no empirical protocol optimization is needed, and that DNA oligonucleotides employed are unmodified and broadly available. The key enabling innovation is a rationally designed competitive hybridization reaction that enables PCR not only to sensitively recognize and selectively amplify even single nucleotide variations (SNVs) at allele frequencies of 0.1%, but also to do so across a temperature window spanning 8*°*C. Our variant allele enrichment method, the Blocker Displacement Amplification (BDA), significantly reduces both the cost and the complexity of profiling rare DNA variants, making genomics analysis more accessible and economical, both for researchers and for clinicians. Compared to other molecular diagnostic technologies used today for detection and quantitation of rare alleles from clinical samples, BDA is unique in simultaneously providing good mutation sensitivity, high mutation multiplexing, fast turnaround, and low reagent and instrument cost (Table 1). Furthermore, in contrast to many proof-of-concept works in academic literature showing high mutation sensitivity against one or a few mutations, we here tested 156 single-nucleotide variants to show the sequence generality of BDA enrichment.

**TABLE 1:** Comparison of rare mutation detection technologies. Innovations to the basic ARMS [12], NGS [15], and microarray [24] technologies have improved mutation sensitivity, but BDA is unique in simultaneously providing good mutation sensitivity, high mutation multiplexing, fast turnaround, and low reagent and instrument cost.

| Method | Mutation LoD | # of Potential Mutations | Protocol Time | Cost per Sample | Instrument Cost |
| --- | --- | --- | --- | --- | --- |
| BDA | 0.1% | 100 - 1000 | 2 hr | $\leq \$3$ | \$650 |
| ARMS | 1%-10% | $\leq 4$ | 2 hr | $\leq \$3$ | \$10,000+ |
| ARMS (castPCR) | 0.1% | $\leq 4$ | 2 hr | $\leq \$3$ | \$10,000+ |
| NGS | 1% | 1000+ | 48+ hr | \$1000+ | \$100,000+ |
| NGS (CAPPseq) | 0.1% | 1000+ | 48+ hr | \$1000+ | \$100,000+ |
| Microarray | 1%-10% | 1000+ | 16+ hr | \$1000+ | \$100,000+ |
| Microarray (MIP) | 0.1%-1% | 1000+ | 48+ hr | \$1000+ | \$100,000+ |

## Blocker Displacement Amplification (BDA) Design

BDA achieves enrichment through enforcing a differential hybridization yield of the forward Primer P to a wildtype (WT) template vs. to a Variant template, resulting in a difference in per-cycle amplification efficiency (Fig. 1). This differential amplification yield is compounded through multiple cycles of PCR to generate high enrichment factors. Hybridization affinity difference is implemented using a Blocker oligonucleotide B whose binding site on the template overlaps with P’s binding site. The simultaneous binding of P and B to the same template molecule is energetically unfavorable; thus, P and B compete in binding through a process of strand displacement [27]. The standard free energy of P displacing B is designed to be marginally positive (thermodynamically unfavorable) in the case of a WT template 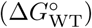, but will be negative for a Variant template due to the mismatch bubble or bulge formed between B and the Variant template.

**FIG. 1:**
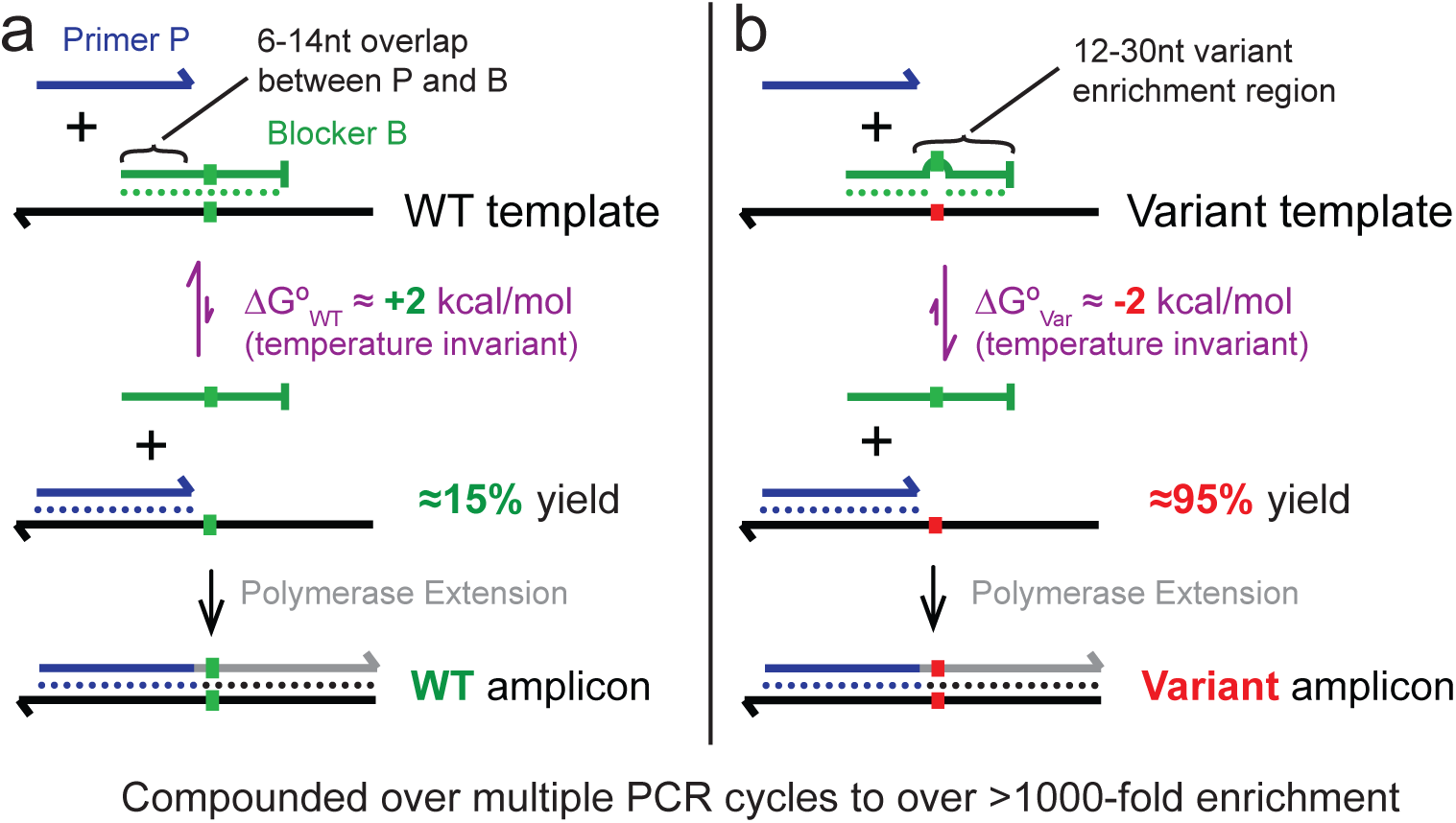
Temperature-robust enrichment of rare alleles by Blocker Displacement Amplification (BDA). **(a)** The Blocker (green) is designed to be perfectly complementary to a known wildtype (WT) template sequence. The 3′ end of the forward Primer sequence (green) is identical to the 5′ end of the Blocker sequence; this 6-14 nt overlap region induces molecular competition between the Primer and Blocker, so that Primer and Blocker hybridization to the same template molecule is mutually exclusive. See Section S1 for Primer and Blocker sequence design considerations. **(b)** The 3′ end of the Blocker contains a 12-30 nt sequence that is not present in the Primer. Any sequence variation in the template in this region will manifest a mismatch bubble or bulge in the Blocker-Template duplex, increasing the favorability of the Primer to displace the Blocker. A single nucleotide variation (SNV) with a characteristic ΔΔ*G°* of 4 kcal/mol [27] would cause the Δ*G°* of the Primer displacing the Blocker on the variant template to drop from +2 kcal/mol to -2 kcal/mol, resulting in approximately 95% hybridization yield at equilibrium. The difference in the Primer displacement favorability results in a difference in per-cycle amplification yield. The amplicon generated by a round of PCR amplification bears the allele of the template, so the amplification yield difference is compounded across many PCR cycles.

The values of 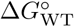 and 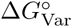 for a given set sequences are only weakly dependent on the temperature of the PCR anneal step, because temperature affects the hybridization stability of both P and B to the template similarly. The competitive hybridization reaction pitting P versus B operates based on a similar principle as the toehold probe [27, 28], which was shown both theoretically and experimentally to exhibit a large affinity difference between SNVs across a large range of temperatures and salinities <u>at equilibrium</u>. However, rapidly changing temperatures and the enzymatic extension of primers in a PCR reaction prevents the solution from attaining equilibrium, necessitating the development of a different, kinetics-driven model for simulating and analyzing BDA behavior. Our model provides design guidelines for maximizing enrichment performance, such as the stoichiometry of B and P. Details of our model are described in Section S2.

In order for BDA to act as an enrichment assay, the sequences of the amplicon must faithfully correspond to the sequences of variant templates originally existing in solution. Consequently, P must bind a conserved sequence upstream of the loci with potential sequence variations (because amplicons always bear the sequence of the primer, and any variations in the primer-binding region of the template would be overwritten). The *enrichment region* of a BDA system thus corresponds to the nucleotides to which the Blocker uniquely binds; templates with sequence variations within the enrichment region are preferentially amplified. The exact length of this enrichment region depends on the sequence in this region, but for the experiments shown here, this length ranges between 12 and 30 nt.

Any of a number of 3′ modifications could be used to effectively prevent the Blocker from being extended by the DNA polymerase during PCR; examples include a minor groove binder [12], an inverted DNA nucleotide, and a 3-carbon spacer [29]. In this manuscript, however, we opted for a simpler approach by adding 4 unmodified nucleotides at the 3′ end of the Blocker that do not pair with the template. For DNA polymerases lacking 3′ to 5′ exonuclease activity, this 3′ “terminator” sequence effectively blocks amplification for all sequences that we tested. We opted for this approach both because such unmodified oligonucleotides are significantly less expensive than functionalized counterparts, and because the thermodynamics of their hybridization are better understood. Although only 2 nt of terminator sequence are needed to suppress Blocker extension, we chose to use 4 nt terminator sequences to account for the possibility of potential Variant templates that happen to match one of the terminator sequence nucleotides, and to account for possible oligonucleotide synthesis errors.

## Results

### Non-pathogenic SNP Results on Genomic DNA

To quantitate the fold-enrichment of BDA on a representative set of SNVs, we first performed enrichment of non-pathogenic SNPs from human genomic DNA (gDNA). NA18537 and NA18562 are genomic DNA samples extracted from two cell lines as part of the 1000 Genomes Project [30], and have well-characterized genotype information for many non-pathogenic SNPs. We generated gDNA samples with a variety of variant allele fractions (VAFs) by mixing the NA18537 and NA18562 samples at different ratios.

Fig. 2abc shows the design and performance of a BDA design for detecting the rs3789806 C>G SNP. The sample with 0.01% VAF can be clearly distinguished from that with 0% VAF, indicating that more than 10,000 × SNV enrichment is achieved. The Sanger sequencing results of amplicons from BDA enrichment of the 0.1% VAF sample (Fig. 2d) verify that BDA functions as an enrichment method, wherein the amplicons reflect the sequences of the original Variant templates. The regression line between the cycle threshold (Ct) and the VAF has slope of 3.45, indicating that PCR amplification of the variant template occurs at near 100% efficiency. This conclusion is supported by the observation that the 100% VAF sample exhibits at similar Ct in PCR reactions with and without the Blocker (see Section S3).

**FIG. 2:**
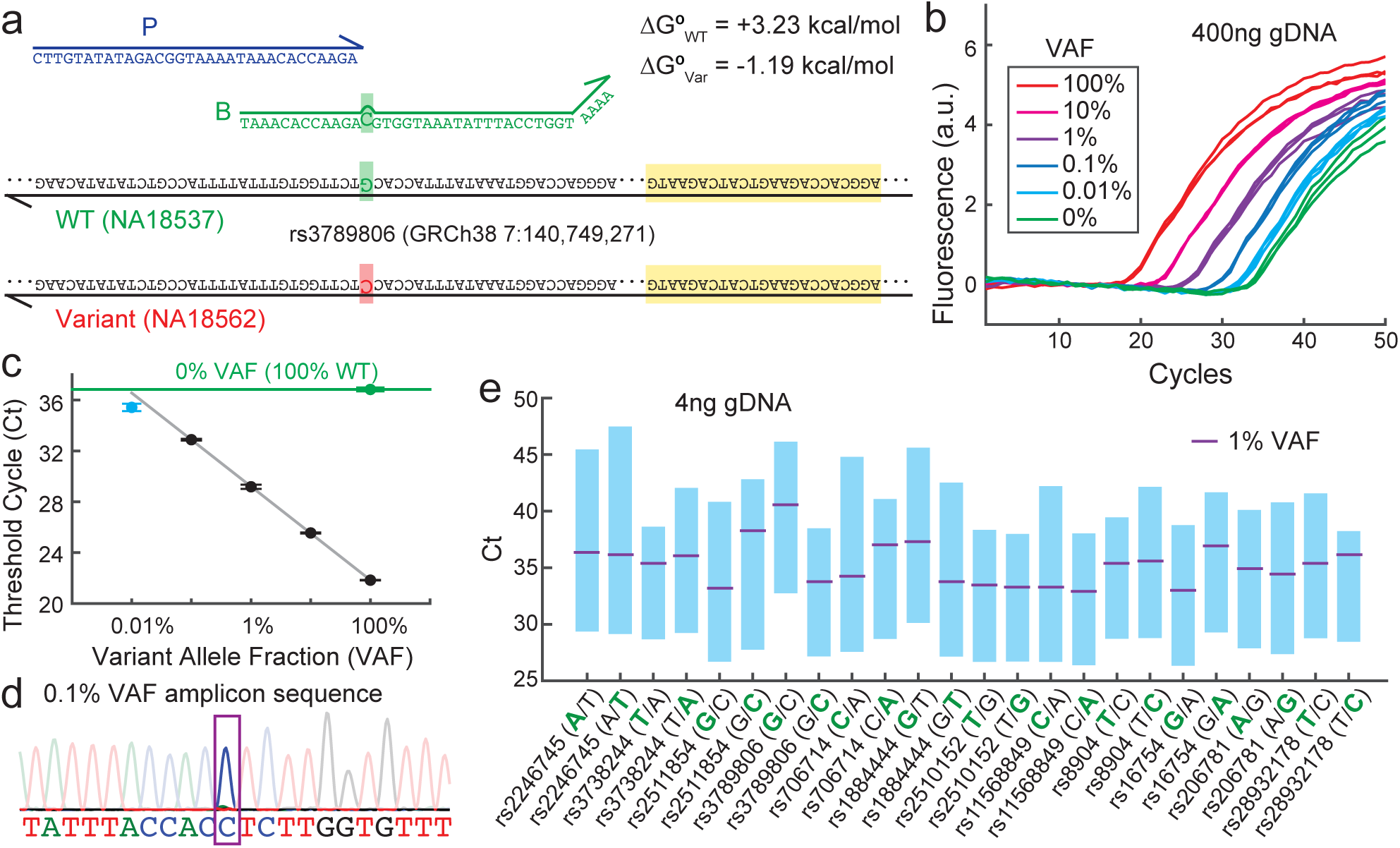
Quantitative PCR (qPCR) results on BDA enrichment. **(a)** Primer and Blocker sequences for enrichment of the non-pathogenic rs3789806 SNP. The reverse primer sequence is highlighted in yellow. The NA18537 genomic DNA (gDNA) sample bears the homozygous WT allele, and the NA18562 gDNA sample bears a homozygous SNP variant. The 4 A’s at the 3′ end of the Blocker sequence serves as a termination sequence to prevent the Blocker from being elongated by the Taq polymerase during PCR. **(b)** Triplicate qPCR results for various mixtures of NA18562 and NA18537 with different variant allele fractions (VAFs). Each sample consisted of 400 ng gDNA in total (corresponding to 120,000 haploid copies), with the indicated VAF fraction as NA18562 and the remainder as NA18537 (e.g. 1% VAF = 4 ng NA18562 and 396 ng NA18537). Experiments were performed using the Applied Biosystems PowerUP supermix using a 2-step protocol alternating 10 s at 95*°*C and 30 s at 60*°*C. **(c)** Summary of threshold cycle (Ct) values for the results observed in panel (b). Error bars show 1 standard deviation; 0.01% VAF data points are not included in linear fit. **(d)** Sanger sequencing of 0.1% VAF amplicon confirms that the Variant allele C is dominant (*>*90% peak height), indicating over 10000-fold enrichment. **(e)** Summary of qPCR results for 24 different BDA designs to 12 non-pathogenic SNPs; both allelic variants of each SNP were enriched using BDA in separate experiments. The bolded green nucleotide indicates the allele to which the Blocker is designed to suppress amplification. The bottom edge of the blue bars indicate the median Ct of the 100% VAF sample, the upper edge of the blue bars indicates the median Ct of the 0% VAF sample, and the purple line indicates the median Ct of the 1% VAF sample. Here, 4 ng gDNA samples were used for each experiment. See Section S3 for raw qPCR traces.

BDA is a sequence-general method that effectively enriches all SNVs regardless of the nucleotide identities of the Vari-ant and WT. Fig. 2e summarizes the qPCR results for 24 separate BDA systems, representing two instances of each of the 12 possible SNV/WT nucleotide pairs; for a large majority of WT/SNV pairs, the median enrichment observed was greater than 1000. The results in Fig. 2e are for preliminary BDA designs that were not subjected to empirical optimization; our later experiments (Fig. 4d) suggest that some Primer/Blocker sequence adjustments may lead to even better enrichment performance.

### Temperature Robustness

An important and novel distinguishing feature of BDA is the broad temperature range over which it effectively enriches variants. Fig. 3ab shows the enrichment performance of the BDA system across anneal/extend step temperatures ranging between 55 and 65*°*C. The Ct values for 1% VAF and 0% VAF samples maintain a difference of more than 10 cycles for anneal/extend temperatures of 56 through 64*°*C. Below 56*°*C, DNA polymerase activity decreases, resulting in poor amplification; above 64*°*C, the hybridization of the primer to either template becomes unstable, also resulting in poor amplification.

**FIG. 3:**
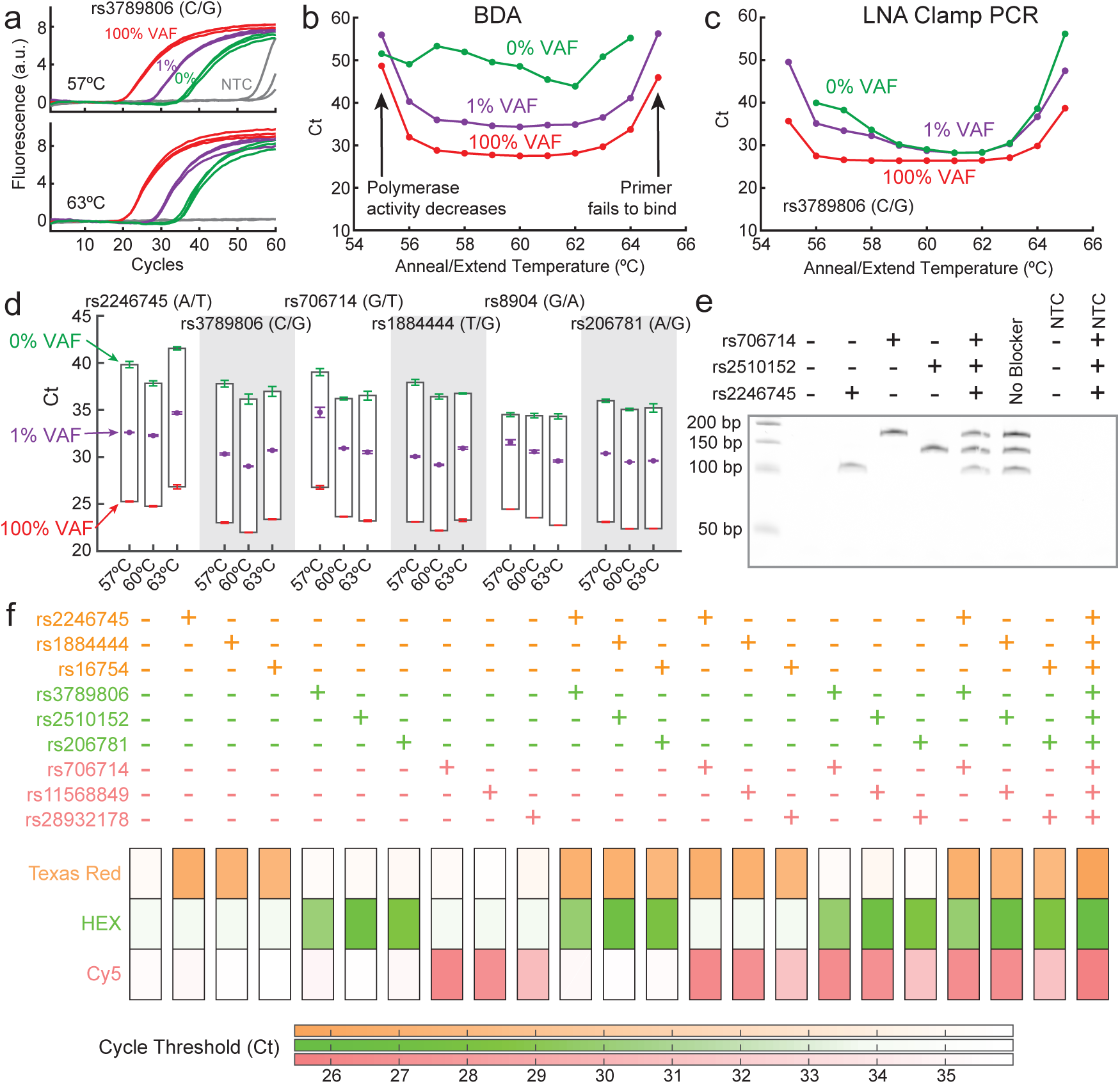
Temperature robustness of BDA. **(a)** Triplicate qPCR traces of 100%, 1%, and 0% VAF samples of the rs3789806 SNP using anneal/extend step temperatures of 57*°*C and 63*°*C; 200 ng gDNA input. **(b)** Summary of Ct values for different anneal/extend temperatures; 4 ng gDNA input. The BDA system efficiently enriches variants across an 8*°*C temperature range, from 56*°*C to 64*°*C. **(c)** Comparative performance of PCR allele enrichment through use of an LNA clamp. Because the binding of the LNA clamp is temperature sensitive, a significant Δ*Ct* is observed only between 56 and 57*°*C. Even at the optimized temperature of 57*°*C, Δ*Ct* values for LNA clamp are lower than BDA. **(d)** Temperature robustness validation for 5 additional SNPs. All tested BDA designs maintain Δ*Ct* ≥ 10 at temperatures from 57 to 63*°*C; 200 ng gDNA input. **(e)** Multiplexed allele enrichment using the portable and inexpensive MiniPCR instrument. Three different BDA systems were designed simultaneously to amplify select alleles of the rs706714, rs2510152, rs2246745 SNPs; amplicons lengths were 171, 138, and 107 nt, respectively, which are clearly distinguished on this 10% denaturing polyacrylamide gel. “-” indicates where the corresponding Blocker suppresses the NA18537 allele, and “+” indicates where the Blocker suppresses an alternative allele. Thus, “+” reactions are designed to provide significant amplification (low Ct). See Section S7 for details. **(f)** Temperature robustness facilitates design of multiplex BDA assays. Here, 9 different BDA systems to the shown SNPs are simultaneously applied to amplify 9 different alleles; each system is reported by a different Taqman probe. Each reaction (vertical column) employs different combinations of blockers to different alleles, but uses the same NA18537 sample. The final concentration of forward and reverse primers are 50 nM each, and the concentration of the Blockers at 500 nM each; 4 ng NA18537 input. See Section S6 for details and qPCR traces.

As a contrast to BDA’s temperature robustness, we also tested enrichment PCR on the rs3789806 SNP using LNA clamp oligonucleotides at a variety of temperatures (Fig. 3c; see Section S5). The LNA clamp provided high amplification yield and significant discrimination between 1% and 0% VAF only at anneal/extend temperatures of 56 to 57*°*C. Our expectation is that similar temperature sensitivities would be observed for other existing PCR enrichment methods [20, 23] [41].

BDA temperature robustness is sequence-generic; Fig. 3d shows the observed Ct values for 6 WT/Variant pairs at 100%, 1%, and 0% VAF. The Ct values of triplicate 100% VAF and 1% VAF samples are generally consistent across all temperatures tested. Some variability observed in the Ct of the 0% VAF sample is believed to be primarily due to the stochastic nature of the early cycles of PCR, specifically in (1)enzyme misincorporation errors and (2) WT amplification yields. These hypotheses are supported by the observation that larger gDNA input quantities result in smaller variability in 0% VAF Ct values; at 4 ng and 200 ng gDNA input, the standard deviation of 96 BDA reactions with 0% VAF were observed to be 4.02 and 0.48 cycles, respectively (Section S4).

The broad temperature robustness of BDA manifests as a strong advantage both in the design and operation of multi-plexed allele enrichment systems, and in the use of portable and inexpensive thermocycling equipment to detect and enrich sequence variants. Fig. 3e shows polyacrylamide gel electrophoresis results of using a 3-plex BDA system on the MiniPCR, an instrument with dimensions of roughly 5 cm × 13 cm × 10cm and a commercially available retail price of $650 [31, 32]. BDA may thus facilitate point of care or field-use nucleic acid testing, for applications such as cancer recurrence monitoring in a primary care physician’s office. The fact that the BDA Primer and Blocker are unmodified DNA oligonucleotides means that reagents are significantly less expensive than modified or functionalized alternatives, rendering BDA desirable for testing in resource-limited settings (e.g. for infectious disease antibiotic resistance diagnostics).

To demonstrate higher multiplexing capability as well as to observe amplification dynamics, we next performed 9-plex BDA on a standard Biorad CFX 96 instrument. The Ct values observed are consistent with expectations based on single-plex results, and the decrease of primer and blocker concentrations necessary for allowing 9-plex amplification does not appear to significantly affect Ct values when cycle times are lengthened to compensate (Fig. 3f). Importantly, the multiplex enrichment experiments summarized in Fig. 3ef represent our first-try designs, and did not undergo any empirical optimization. We believe that significantly higher multiplexed allele enrichment can be achieved with BDA; however, methods of readout more sensitive and multiplexed than traditional qPCR instruments are needed to fully capture the value of multiplexed BDA.

### Model vs. Experiments

We model the BDA reaction in two parts: (1) a mass-action kinetic simulation of the hybridization and enzyme elongation reactions at each anneal/extend step to calculate the per-step amplification yield for the WT and Variant templates, and (2) a discrete simulation of each PCR cycle using the amplification yields from the previous step. Where possible, we used literature-reported rate constants and standard free energies; two parameter values were estimated *ad hoc*. The purpose of this model is to understand the effects of various design parameters on the enrichment performance of BDA; accurate quantitative match between simulation and experimental traces is secondary (Fig. 4a). Deviations between model and experimental results likely reflect both the simplicity of the model and the inaccuracy of literature reported thermodynamics parameters [25, 33].

**FIG. 4:**
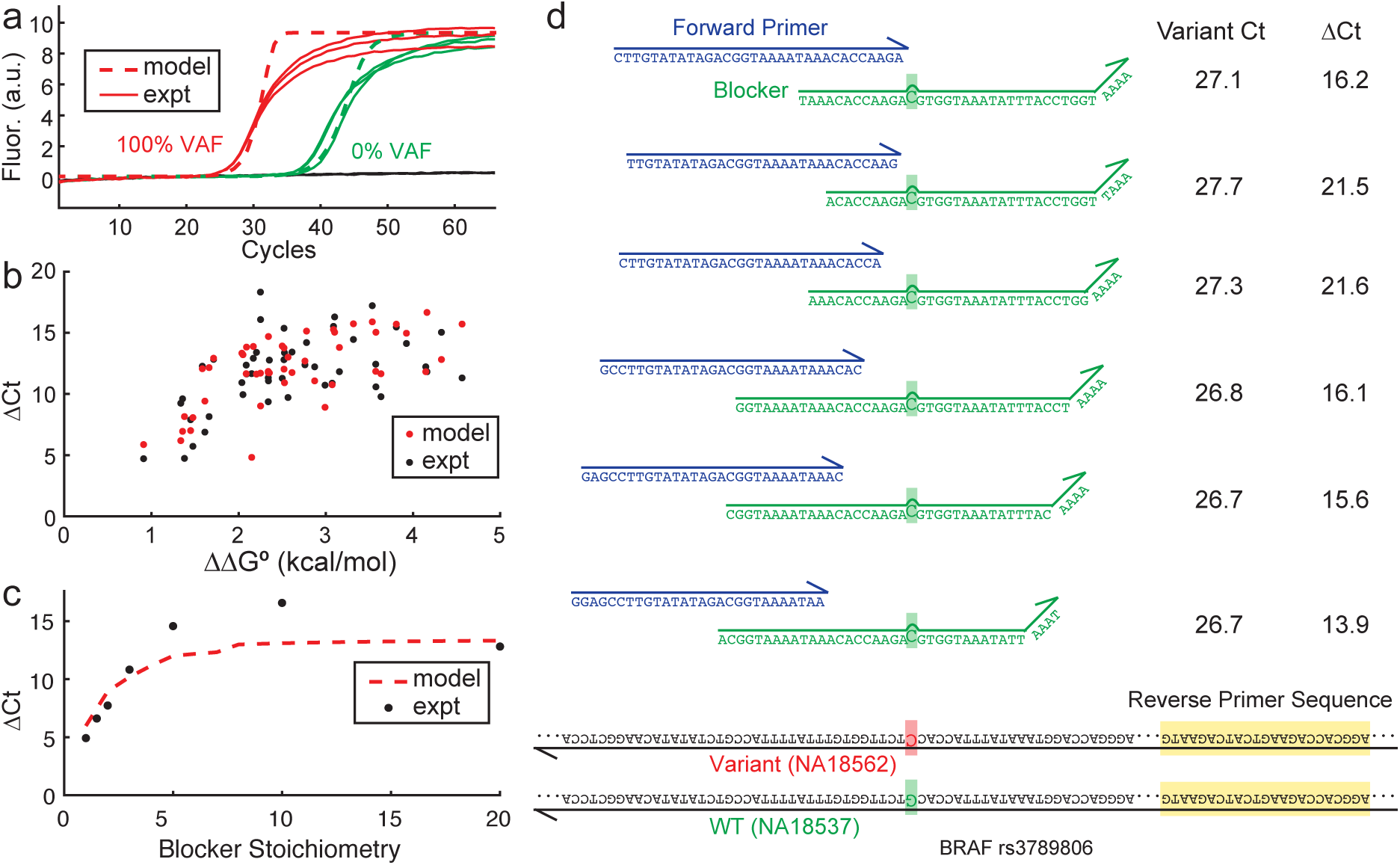
Model guidance of BDA sequence and protocol design. **(a)** Our model of the BDA reaction process captures many aspects of the BDA, and furthermore suggests optimal conditions under which BDA provides the best enrichment. Here, we show a comparison of experimental (solid lines) and simulated BDA qPCR traces (dashed lines) for BDA enrichment of the rs2246745 SNP. See Section S2 for model and fitted parameter details. **(b)** Model (red dots) and experimental (black dots) effects of mismatch ΔΔ*G°* for all SNPs tested in Fig. 2e. Variants with more destabilizing thermodynamics (large ΔΔ*G°*) yield larger ΔCt, up to about 2.5 kcal/mol, after which ΔCt plateaus. **(c)** Model (dashed line) and experimental (black dots) effects of Blocker stoichiometry for BDA on the rs1884444 SNP. Blocker stoichiometry is defined as the initial concentration of the Blocker divided by the initial concentration of the forward Primer. Maximal Ct is achieved when Blocker stoichiometry ≥7. To be conservative against slight model errors, we chose Blocker stoichiometry of 10 for all experiments in this manuscript other than the ones shown in this panel. **(d)** Our model predicts many BDA systems to yield similar ΔCt for a given variant. All six designs tested here yielded ΔCt *>* 13 with 4 ng gDNA input.

Our model suggests that ΔΔ*G°* is a key parameter that sets the upper limit for the achievable enrichment by BDA. ΔΔ*G°* denotes the difference in the standard free energy of hybridization of the Blocker binding to the WT as compared to binding to the Variant, and is primarily dependent on the sequence identities of the WT and Variant alleles. At 60*°*C and typical PCR buffer conditions, ΔΔ*G°* values range between 0.8 kcal/mol and 5 kcal/mol. The model suggests that larger ΔΔ*G°* results in larger ΔCt values, with saturation at roughly 3 kcal/mol. Thus, some Variant/WT pairs with smaller ΔΔ*G°* will exhibit lower enrichment by BDA due to their sequence identities.

For enrichment of a specific Variant sequence, many BDA designs will yield good enrichment, as long as design principles are followed (Fig. 4d). This flexibility further facilitates the design of highly multiplexed BDA systems, because specific BDA designs for enriching different loci may be incompatible with one another (e.g. tendency to form primer dimers).

### Hotspot Multiplexing

BDA enriches all Variants with sequence differing from the WT in the enrichment region. For a typical 20 nt enrichment region, 60 SNVs can be simultaneously enriched by a single BDA system, albeit with differing fold-enrichment because of varying ΔΔ*G°* values. In this regard, BDA and other enrichment PCR techniques are significantly superior to allele-specific PCR (e.g. ARMS) approaches — each plex of BDA is the equivalent of a 60-plex ARMS. To rigorously demonstrate that BDA does enrich all potential variants in the enrichment region, we systematically designed 2 sets of synthetic DNA templates bearing SNVs, corresponding to all SNVs in a 18 nt enrichment region on chromosome 5 and a 21 nt enrichment region on chromosome 11 (Fig. 5a). The experimentally observed distribution of ΔCt values for these SNVs range between 7 and 14, with mean and median both at roughly 10.5 (corresponding to over 1000-fold enrichment). Note that the same BDA primer/blocker system was used to enrich all 54 SNVs in the left panel, and another BDA system was used to enrich all 63 SNVs in the right panel. In applications where BDA is used to detect/quantitate a suspected known mutation (Fig. 2e), significantly higher ΔCt is achieved because the BDA system can be designed to either the forward strand or the reverse strand, depending on which exhibits the larger ΔΔ*G°* value (Fig. 4b, see also Section S8).

**FIG. 5:**
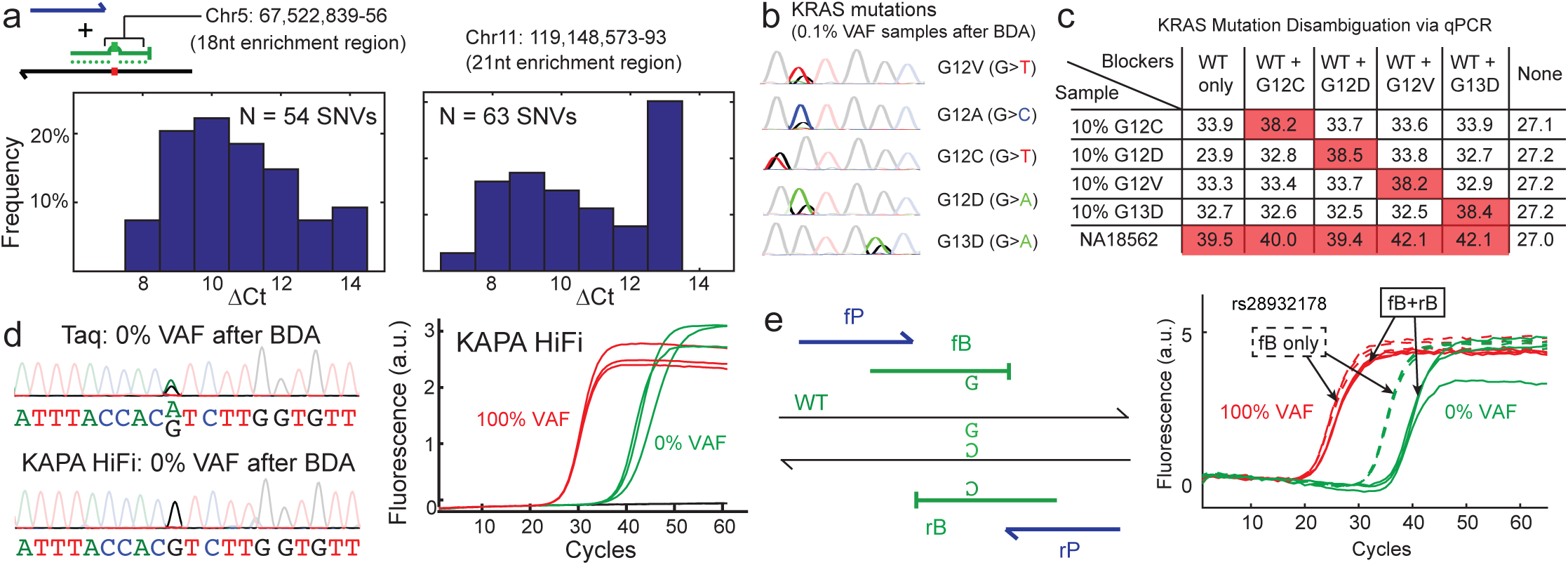
Hotspot multiplexing and improving mutation sensitivity. **(a)** BDA preferentially amplifies all variants within the enrichment region. The left panel shows the distribution of experimental ΔCt values observed for the same BDA primer/blocker set to 54 synthetic templates each bearing a SNV within a 18 nt enrichment region, and the right panel shows the ΔCt value distribution for 63 synthetic SNV templates for a different BDA system. See Section S8 for qPCR traces; Ct values are median of triplicates. **(b)** A single-plex BDA to enrich 0.1% VAF of 5 KRAS mutants from cell line genomic DNA; Sanger sequencing results of amplicon products are shown. **(c)** Variant disambiguation in qPCR setting using Blocker specific to the Variant of interest. The simultaneous use of a WT Blocker and a Mutant Blocker produces weak amplification (high Ct) when no incidental SNPs are present, but strong amplification (low Ct) if one or more incidental SNPs are present. Ct values shown are median of triplicates. **(d)** BDA using the Taq polymerase is limited by enzyme misincorporation events. In the enrichment region of the BDA system amplifying a purely WT sample for rs3789806, G*>*A misincorporation events (boxed) resulted in a significant fraction of the amplicons bearing variant sequence. Similar BDA on pure WT sample using the KAPA high-fidelity DNA polymerase does not result in visible *de novo* variations. Because KAPA HiFi possesses 3′ to 5′ exonuclease activity, we used a 3-carbon spacer at the 3′ end of B to prevent extension. **(e)** Improved variant enrichment using dual BDA to block WT amplification on both the forward and reverse template strands. fB and rB denote forward Blocker and reverse Blocker that interact with the forward Primer fP and reverse Primer rP, respectively. For this (particularly challenging) Variant/WT pair, the ΔCt between 100% Variant and 100% WT is improved from 8.6 (single BDA) to 15.0 (dual BDA), using 400 ng gDNA input.

Hotspot multiplexing is important for a variety of molecular diagnostics applications. For example, codons 12 and 13 in the KRAS oncogene is a hotspot region in which 12 different non-synonymous point mutations have been observed. Mutation status of KRAS can information clinical treatment; colorectal cancer patients are resistant and not recommended to be prescribed cetuximab.

We designed a BDA system to query variants in codons 12 and 13 of the KRAS gene. To test the BDA system, we diluted 5 reference cell line gDNA samples with different KRAS mutations (Horizon Discovery) with WT cell line DNA to generate 0.1% VAF samples. Sanger sequencing of the amplicons after BDA (Fig. 5b) confirms that amplicon sequence identity matches that of the original sample mutations. To unambiguously detect a specific Variant of interest, two separate BDA reactions should be run: one with a standard WT Blocker, and one containing both the WT Blocker and a Blocker to the Variant of interest (Fig. 5c). Significant amplification (low Ct) of a sample with the second BDA reaction indicates that the sample contains an incidental variant other than the Variant of interest.

### Enhancing Enrichment

The allele enrichment achieved by BDA is limited by two factors: (1) DNA polymerase nucleotide misincorporation events that create *de novo* Variants subsequently enriched by BDA [35], and (2) imperfect suppression of WT template amplification by the Blocker. Limitation (1) can be mitigated through the use of highfidelity DNA polymerases with 3′ to 5′ proofreading activity (Fig. 5d), and (2) can be mitigated through the use of a second Blocker for the reverse primer (Fig. 5e). For different Variant/WT pairs, either (1) or (2) may be dominant; thus, addressing each of these two problems results in improved BDA performance for a subset of Variant/WT sequence pairs. Judicious application of these two approaches allows BDA enrichment to be enhanced for a large fraction of, if not all, Variant/WT pairs.

### Clinical Cell-Free DNA Analysis

Cell-free DNA (cfDNA) consists of short (approx. 160 nt) double-stranded DNA molecules present in blood plasma at roughly 5 to 20 ng/mL [1]. Cancer-specific mutations in cfDNA can be used for noninvasive therapy guidance or recurrence monitoring [2, 3, 38], and even holds potential as cancer early detection biomarkers [39]. Fig. 6 shows the BDA qPCR results for the analysis of cell-free DNA from the blood plasma of 7 lung cancer patients, collected between Jan 2013 and Nov 2014 (see also Section S9). Informed consent was obtained from all subjects. As a comparison, the cfDNA mutations were also profiled by deep sequencing using molecular lineage tags to suppress sequencing and PCR errors via a modified version of the technique presented in ref. [36]. There is general agreement on mutant VAF between the BDA qPCR and the deep sequencing methods; differences observed may be due to unequal distribution of mutant cfDNA during sample splitting (sample 2), or due to differences in mutation VAF based on different length cfDNA fragments (sample 5).

**FIG. 6:**
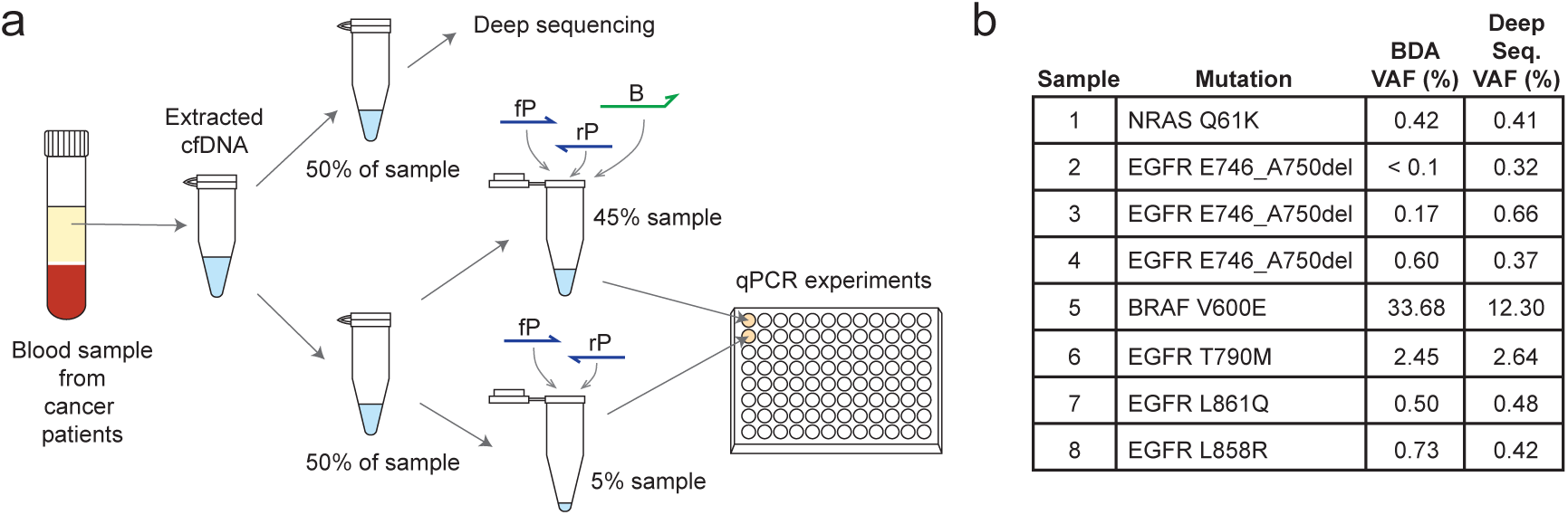
Detection of somatic cancer mutations in clinical cell-free DNA samples extracted from blood plasma. **(a)** Pre-analytic workflow. **(b)** Inferred mutation VAF of clinical samples using BDA, as compared to using deep sequencing [36, 37]. The observed difference in some samples between deep sequencing and BDA may be due to unequal partitioning of rare mutation-bearing cfDNA molecules at near the Poisson limit.

For example, tumor-derived cfDNA fragments are typically shorter than normal cfDNA, so the choice of amplicon length may impact the observed allele frequency.

## Discussion

BDA overcomes the temperature sensitivity that limit all existing PCR enrichment methods through a rationally designed competitive hybridization reaction. This temperature robustness translates into practical advantages for molecular diagnostics and genomics research in at least two ways: facilitating the highly multiplexed enrichment of many SNVs, and enabling the use of more portable and economical PCR instruments with lower temperature accuracy and uniformity. BDA does not rely on expensive functionalized DNA oligonucleotides, is broadly compatible with many enzymes, and is tolerant to instrument temperature inaccuracies and non-uniformity; consequently, BDA is highly economical and accessible to clinicians and researchers. Systematic experimental validation of BDA enrichment on 156 different SNVs, and on a variety of samples ranging from synthetic templates to cell line genomic DNA to clinical blood samples, shows the robustness of BDA as a rare variant enrichment and detection method.

An immediate area of clinical application for BDA technology is noninvasive cancer recurrence monitoring based on cfDNA. NGS-based solutions for analyzing cfDNA DNA are too costly to be repeatedly and regularly applied, while low-multiplex PCR methods are insufficient for high clinical sensitivity. In this manuscript, we demonstrated multiplex BDA targeting 9 different genomic regions within the same reaction; simultaneously, we showed that each BDA system is capable of enriching and detection 50+ variants. Together, these indicate that roughly 500 different SNVs can be detected at roughly 0.1% allelic frequency in a single low-cost and rapid reaction. Additional development and optimization of the BDA technique could potentially further extend the number of mutations detected and enriched to over 1000, suggesting a rare variant discovery as another potential application for BDA.

The fold-enrichment achieved for SNVs using the basic BDA implementation (lacking high fidelity enzyme and dual blocking) was observed to be between approximately 300 and 30,000, varying based on SNV and WT sequences. This performance could be improved through empirical optimization, as the DNA hybridization thermodynamics parameters we used for designing BDA systems are known to be imperfect [25]. In settings where open-tube procedures are allowed (e.g. NGS library preparation), multiple rounds of BDA enrichment followed by dilution could result in theoretically unlimited rare mutation sensitivity.

## Acknowledgements

The authors thank Jin H. Bae and J. Sherry Wang for experimental advice. The authors thank Dmitriy Khodakov for initial testing of BDA on the MiniPCR platform, Ping Song for assistance with gel electrophoresis, and Azeet Narayan for testing BDA in the Patel lab. This work was funded by CPRIT grant RP140132 to DYZ and NIH grant R01CA203964 to DYZ. LRW conceived the project, designed and conducted the experiments, analyzed the data, and wrote the paper. SXC designed the experiments and analyzed the data. YW conducted the experiments and analyzed the data. AAP provided clinical samples and analyzed the data. DYZ conceived the project, guided experimental design, analyzed the data, and wrote the paper. Correspondence may be addressed to D.Y.Z. There is a patent pending on the BDA system described in this work.

